# The Effects of Incline Level on Optimized Lower-Limb Exoskeleton Assistance

**DOI:** 10.1101/2021.09.13.460170

**Authors:** Patrick W. Franks, Gwendolyn M. Bryan, Ricardo Reyes, Meghan P. O’Donovan, Karen N. Gregorczyk, Steven H. Collins

**Affiliations:** Department of Mechanical Engineering, Stanford University, Stanford, CA, 94305 USA; DEVCOM Soldier Center

**Keywords:** Exoskeleton, walking assistance, human-in-the-loop optimization, metabolic cost, incline

## Abstract

For exoskeletons to be successful in real-world settings, they will need to be effective across a variety of terrains, including on inclines. While some single-joint exoskeletons have assisted incline walking, recent successes in level-ground assistance suggest that greater improvements may be possible by optimizing assistance of the whole leg. To understand how exoskeleton assistance should change with incline, we used human-in-the-loop optimization to find whole-leg exoskeleton assistance torques that minimized metabolic cost on a range of grades. We optimized assistance for three expert, able-bodied participants on 5 degree, 10 degree and 15 degree inclines using a hip-knee-ankle exoskeleton emulator. For all assisted conditions, the cost of transport was reduced by at least 50% relative to walking in the device with no assistance, a large improvement to walking that is comparable to the benefits of whole-leg assistance on level-ground. This corresponds to large absolute reductions in metabolic cost, with the most strenuous conditions reduced by 4.9 W/kg, more than twice the entire energy cost of level walking. Optimized extension torque magnitudes and exoskeleton power increased with incline, with hip extension, knee extension and ankle plantarflexion often growing as large as allowed by comfort-based limits. Applied powers on steep inclines were double the powers applied during level-ground walking, indicating that larger exoskeleton power may be optimal in scenarios where biological powers and costs are higher. Future exoskeleton devices can be expected to deliver large improvements in walking performance across a range of inclines, if they have sufficient torque and power capabilities.

## I. Introduction

**E**XOSKELETONS should be able to assist walking on inclines if they are to be effective in real-world settings. Incline walking is associated with increased fatigue and decreased walking speed [1], and exoskeleton assistance is well-suited to reduce these negative outcomes. A number of exoskeletons to date have assisted walking by reducing the metabolic cost of walking [2], [3], [4], [5], [6], [7], [8], [9], [10], [11], [12], [13], [14], which has been identified as an important outcome metric for evaluating exoskeletons [2], [15]. While these improvements are promising, most devices have focused on assisting level-ground walking [2]. If exoskeletons could assist incline walking as well, they could could deliver large improvements to functional mobility outcomes, because steeper inclines result in higher metabolic energy costs and are more fatiguing than level ground [16], [17], [18]. Assisting incline walking could improve maximum performance for military personnel, first responders, or other users who traverse hills, mountains, or other difficult terrain [19], [20]. In urban environments, incline assistance could help older adults or people with impairments navigate ramps, stairs, and hills, where loss of strength and increased fatigue can be a limiting factor [21], [22], [23], [24], [25], [26].

Some exoskeletons have assisted walking on inclines, but larger improvements may be possible. Single-joint exoskeletons have reduced the metabolic cost of walking up inclines [13], [27], [28], [29], with the largest reductions around 15.5% relative to no exoskeleton using a hip-only device [29]. This improvement is slightly smaller than single-joint assistance on level-ground (around 18% relative to walking in no exoskeleton) [9], [12]. It is possible that exoskeletons might deliver larger improvements to incline walking than level-ground walking, because steeper inclines require larger biological joint powers [30], [31], [32] and incur higher metabolic energy costs [16], [17], [18], giving a larger opportunity for the device to effectively assist. Alternatively, it is possible that assisting incline walking may be less effective because of the related effect of walking speed: people tend to walk slower up inclines [33], [34], [35], [36], and exoskeletons have been less effective at slower speeds [7], [37]. To see if larger improvements to incline walking are possible, the best strategy may be to optimize whole-leg assistance using a device with large torque capabilities [7], [38].

Assisting the whole leg during incline walking could lead to greater reductions in metabolic cost. Single-joint devices have focused primarily on the ankle and hip for level-ground walking, where they do most of the biological work of the leg [30], [31]. Each of the devices that have reduced the metabolic cost of walking on inclines have also assisted only a single joint [13], [27], [28], [29], [39]. However, the hips, knees, and ankles all contribute to the total positive power of the leg during incline walking [30], [31], so assisting all joints simultaneously could be best. While knee assistance has been the least effective of all joints on level ground [38], biological knee torque and power increase as grade increases [31], meaning the knee may be increasingly helpful to assist with inclines. For level-ground walking, whole-leg assistance produced greater improvements to metabolic cost compared to single-joint and two-joint assistance [38], consistent with expectations from biomechanical simulations [40], [41]. It is likely that whole-leg assistance could deliver maximum benefits to walking up inclines as well.

By understanding how optimal whole-leg assistance changes with incline, exoskeleton designers could build mobile devices that are most effective in real-world settings. While biological torques change with incline, including large increases in hip extension torque and smaller increases in knee extension and ankle plantarflexion torques [31], it is unknown how exoskeleton torques should change with incline as well. If we understood the relationship between incline level and optimal assistance parameters, mobile devices could adapt their control strategy online to best assist current terrain. In addition, understanding this relationship could specify the needed torque and power capabilities for future exoskeleton products. This is important for mobile device design, since actuation capabilities often come in direct tradeoff with device mass and cost. Finally, knowing how metabolic reductions correspond to these assistance parameters could set expectations for what mobile devices could achieve.

The purpose of this study was to identify how changing incline affected optimized whole-leg exoskeleton assistance and the corresponding reductions in metabolic cost. We hypothesized that exoskeleton assistance may produce greater metabolic reductions with increasing inclines. Since biological joint power increases as incline increases, we expected that optimal exoskeleton torques and powers would increase as well. We tested these ideas using human-in-the-loop optimization [7] with a hip-knee-ankle exoskeleton emulator [42] that was previously successful in optimizing whole-leg assistance to reduce metabolic cost [37], [38]. Three able-bodied, expert participants walked on a treadmill at 0, 5, 10, and 15 degrees. For each incline level, we used human-in-the-loop optimization to find the applied whole-leg exoskeleton torques that minimized the measured metabolic cost of walking. We measured the metabolic cost of walking and cost of transport to evaluate the effect of incline. We studied how optimized torques and powers changed with incline and how the user’s joint kinematics and muscle activities changed with assistance. By finding and comparing optimized assistance at a variety of inclines, we expected these results to lead to the design of more effective exoskeletons.

## II. Methods

We used human-in-the-loop optimization to find whole-leg assistance that minimized metabolic cost for three incline-walking conditions. Users walked in a hip-knee-ankle exoskeleton (Fig. 1) [42] on an incline-adjustable split-belt treadmill. Assistance was optimized for walking on a 5 degree incline at 1.25 m/s, on a 10 degree incline at 1.00 m/s, and a 15 degree incline at 0.75 m/s. We optimized each incline condition in increasing order of slope, initializing assistance for each new condition with the optimum of the previous condition for each participant. We completed a validation experiment after each optimization to evaluate the effects of incline level and assistance. For each slope, walking in the exoskeleton with assistance was compared to walking in the exoskeleton without assistance, and walking without the exoskeleton. The effect of incline was also assessed by comparing to walking conditions on level-ground at 1.25 m/s, which were previously collected for these participants in this device [38].

**Fig. 1.**
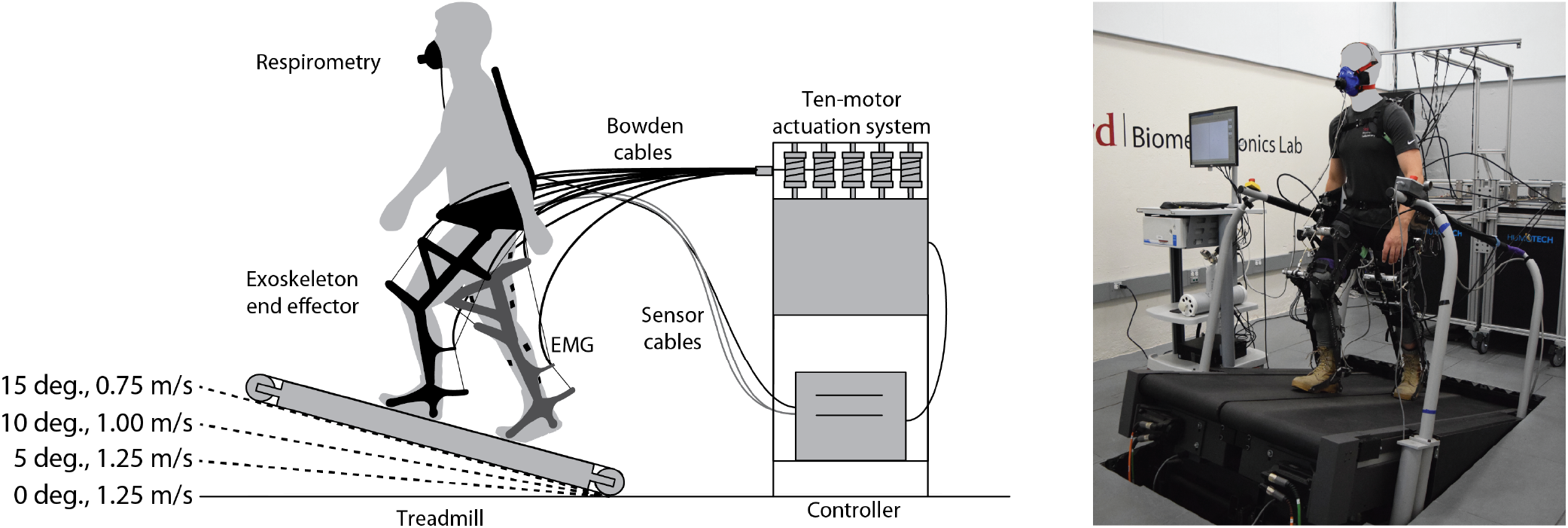
Experimental Setup. (Left) Overview of exoskeleton emulator. Ten powerful off-board motors actuate a lightweight end effector worn by a user who walks on a split-belt treadmill. Metabolic cost is measured using a respirometry system and muscle activity is measured using electromyography (EMG). The tested inclines and corresponding walking speeds are shown. (Right) Photo of experimental setup with user walking on the treadmill at a 15 degree incline, the maximum of the range studied. For reference, an ADA-compliant ramp has a maximum incline of 5 degrees. Participant gave permission for use of their image.

### A. IRB Approval

All user experiments were approved by the Stanford University Institutional Review Board and the US Army Medical Research and Materiel Command (USAMRMC) Office of Research Protections as eProtocol 46935 on 01/09/2019. All participants provided written informed consent before their participation as required by the approved protocol.

### B. Participants

Three healthy participants were included in this study (P1: M, 26 years old, 90 kg, 187 cm; P2: F, 27 years old, 61 kg, 170 cm; P3: M, 20 years old, 84 kg, 176 cm). All three participants were experienced with the device, having recently completed optimization of whole-leg assistance for level ground at 1.25 m/s [38]. These trained participants were expected to quickly adapt to new walking conditions and to new assistance patterns. These participants were also authors of the study (PF, GB, RR) as these were the people who were trained exoskeleton users and had the time to participate in this study.

We were limited to three participants because of the extensive time required to complete the protocol and the difficulty of recruiting participants in the COVID-19 pandemic. Each participant completed at least 50 hours of experiments for this study as part of a longer optimization protocol of individual joints, walking speeds, and loads, totaling between 150 and 300 hours of experiments for each participant. With three participants, we have a statistical power of 0.8 to detect metabolic cost reductions greater than 24%, assuming metabolic cost reductions have a standard deviation of 7.4% [7], meaning we can confidently detect the large reductions we expected [38].

### C. Exoskeleton hardware

Assistance was applied using a hip-knee-ankle exoskeleton emulator (Fig. 1) [42]. The exoskeleton emulator has ten off-board motors connected to a worn end effector through a Bowden cable transmission. The device is capable of applying at least 200 Nm of torque and 2.8 kW of power at each joint. The end-effector has a worn mass of 13.5 kg. Each participant had been previously fit to the device which is adjustable in length at the thigh and shank, adjustable in width at the hips, thighs, and knees, and can be equipped with boots of different sizes.

### D. Exoskeleton control

The exoskeleton is controlled by applying torques at each joint [42]. When no torque is desired, the transmission is slack to ensure no interference with the user. We define control of the joint torques as a function of percent stride. We calculate percent stride using heel strikes detected from ground reaction forces, with the average stride time being the rolling average over the past twenty strides.

For the hips and ankles, the desired torque profile is a spline (piecewise cubic hermite interpolating polynomial) defined by nodes. For the knees, torque is controlled both by percent stride and by state-based control [38]. Knee torque is first defined as a virtual spring during stance, with torque proportional to knee angle. In late stance and around push-off, knee torque is defined as a spline as a function of percent stride. Finally, in late swing, knee flexion torque is commanded as a virtual damper, with torque proportional to knee velocity. The onset and offset of the state-based knee controllers is defined by percent stride.

The desired torques were set using 22 parameters in total that could be adjusted by the optimization algorithm (A-A). The profiles were defined in the same way as previous optimizations of assistance with this device [37], [38]. For the hips, eight parameters were used to define the peak torque magnitude, peak torque timing, and rate of rise and fall of torque, for both hip flexion and hip extension. For the knees, ten parameters defined the stiffness of the virtual spring, the magnitude of flexion torque near push-off, the damping coefficient during late stance, and the onset and offset of each of these torques. For the ankles, four parameters defined the peak ankle torque, the timing of peak torque, and the rise and fall of torque, similar to previously successful ankle exoskeleton optimizations [7]. Each parameter had minimum- and maximum-allowed values based on user comfort during pilot tests and previous experiments. In a few cases, the optimization converged at the limit for a parameter. For the hips, maximum hip extension torque magnitude was set to 0.9 Nm/kg, which was reached during optimization of walking at 10 and 15 degrees. For the ankles, peak ankle torque was set to 0.8 Nm/kg for level-ground walking, and increased to 0.9 Nm/kg for incline walking, both of which were reached during optimization. For the ankles, peak plantarflexion timing was set to be as late as 55% of stride, which was converged upon in a number of conditions, similar to previous optimizations of ankle assistance [7], [37], [38]. These comfort-based parameter limits were first set based on previous experiments with this device [38]. Torque magnitude limits were then updated for incline walking during pilot testing at 5 degrees by increasing torque magnitudes until the user stated torques were too large to walk in comfortably.

To track desired joint torques, we used a combination of closed-loop proportional control, joint-velocity-based compensation, and iterative learning [42], [43]. This approach is accurate in applying torques, with root-mean-square (RMS) errors of 0.79 Nm at the hips and 0.36 Nm at the ankles during whole-leg assistance. Error is typically highest at the knees because there can be discrete jumps in desired torque at the onset and offset of the state-based controllers. For the knees during level-ground walking, RMS error was 2.84 Nm. For higher inclines, the virtual knee extension spring in stance optimized to maximum stiffness, which led to instantaneous increases to large desired torques that were impossible for the emulator to track, resulting in larger RMS errors of 11.05 Nm on average. This tracking error was expected, and our analysis of exoskeleton torques focuses on the measured torques applied to the user. During periods of no torque, the transmission was effectively slack, with an RMS applied torque of less than 0.61 Nm at the hips, 1.13 Nm at the knees, and 0.26 Nm at the ankles.

### E. Optimization Protocol

We optimized assistance first at 5 degrees at 1.25 m/s, then at 10 degrees at 1.0 m/s, and finally at 15 degrees at 0.75 m/s. Slower speeds were chosen for steeper inclines to limit user fatigue and avoid anaerobic respiration, which would disrupt the metabolics estimates. To choose the speeds, we conducted a pilot test where a participant walked in the exoskeleton without torque at each incline, and speed was decreased from 1.25 m/s if needed until the respiratory quotient was less than one. Varying the speed also improves functional relevance, since people walk at slower speeds for steeper inclines [33], [34], [35], [36].

We used human-in-the-loop optimization to find the best exoskeleton assistance for each incline condition. This approach has been previously successful for this device [37], [38] as well as for other exoskeletons [7], [9], [44], [45]. We used covariance matrix algorithm-evolutionary strategy (CMA-ES) [46] to optimize assistance. The goal of the optimization was to minimize measured metabolic cost, which was estimated for each condition after two minutes of walking using a firstorder dynamical model [47], similar to previous work [7], [37], [38]. To attempt to speed convergence, the initial parameter values for each optimization were based on the participant’s optimized assistance from the previous condition (e.g., P2’s optimized assistance at 5 degrees was used as the initial seed for P2’s optimization at 10 degrees).

The duration of optimization was chosen based on previous experiments to be long enough to ensure convergence, while being short enough to be experimentally feasible [38]. This experiment was conducted with experienced users who are likely to adapt quickly to novel walking conditions and new assistance strategies compared to novice users. Initial conditions chosen from previously optimized assistance should also speed convergence. Optimization for each condition consisted of nine generations over at least three days, with each generation requiring 26 minutes of walking. Participants were permitted but not required to take breaks between generations. Participants were permitted to wear wireless headphones to listen to podcasts while walking.

### F. Validation Protocol

A validation experiment was performed after each optimization to accurately assess the effect of incline on walking and the effect of exoskeleton assistance, similar to [37], [38]. During each validation experiment we measured metabolic cost, applied torque, kinematics, applied power, muscle activity, and stride frequency. Each incline level was assessed on a separate day, following optimization of that level and prior to beginning optimization of the next incline level. Bouts of walking in the exoskeleton conditions were longer than the control conditions to ensure that users were adapted to the device and to the assistance, ensuring accurate measurements of steady-state metabolics and of walking strategy. For each validation experiments, users stood quietly for 6 minutes, walked without the exoskeleton for 6 minutes, walked in the exoskeleton with no torque applied for 10 minutes, and walked with the optimized assistance torques for 20 minutes, in a double-reversed order (ABCDDCBA). We did not randomize this order to ensure maximum acclamation to the device for the exoskeleton assistance conditions, and because of the time it takes to don and doff the device. For the no exoskeleton condition, users wore the same type of boots that are included in the exoskeleton. Users were required to rest for at least three minutes between walking conditions, and for at least five minutes before quiet standing, to ensure their metabolics returned to baseline.

### G. Measured Outcomes

We collected biomechanical data of the user when walking in the different conditions during the validation. We calculated the average of these measurements over the last three to five minutes of walking of each condition to ensure the user’s metabolics and gait had reached steady-state. All conditions were evaluated twice and measurements were averaged across both evaluations.

#### 1) Metabolic Cost and Cost of Transport

We calculated metabolic cost using indirect calorimetry. We measured oxygen consumption (VO2), carbon dioxide expulsion (VCO2), and breath duration on a breath-by-breath basis (Quark CPET, COSMED). Metabolic rate was calculated for each condition using a modified Brockway equation [7]. For each incline level, metabolic cost was calculated during quiet standing, walking with no exoskeleton, walking in the exoskeleton without assistance, and walking with assistance. To calculate the cost of walking, the cost of quiet standing was subtracted from the walking conditions. To ensure accurate metabolic measurements, users fasted for at least two hours before optimization and at least four hours before validations. To evaluate changes in metabolic cost, we used two-tailed paired t-tests to calculate statistical significance.

For all validations of incline conditions, the participant wore a cloth mask underneath the metabolics mask to comply with COVID-19 safety protocols. Level-ground optimizations and validations were performed without cloth masks prior to the pandemic [38]. Participants 2 and 3 wore a cloth mask for all incline optimizations and validations. Participant 1 wore a cloth mask for all incline validations and for optimizations at 10 and 15 degrees. Optimization of participant 1 walking at 5 degrees occurred prior to needing cloth masks, but validation for walking at 5 degrees was re-evaluated with a cloth mask (4 months later) to ensure all incline conditions were evaluated similarly. The presence of the cloth mask did affect the metabolic measurements, by creating a downward offset in the measured metabolic cost [38] (A-B). This may mean slightly underestimating the absolute measurements of metabolic cost of incline conditions, but we expect changes between conditions to be consistent and for percent reductions in metabolic cost to be accurate, because we are comparing masked conditions. It is possible that the mask effect is non-linear with rate of breathing, meaning it is possible that we underestimate more strenuous conditions more so than less-strenuous conditions.

We calculated the cost of transport by dividing the metabolic cost of walking by the walking speed. Walking speed decreased with incline, and evaluating the cost of transport minimized the effect of speed. While speeds were selected to keep the respiratory quotient below 1 for accurate estimation of metabolic rate, there were short periods of respiratory quotients at or above 1 for two subjects during the steepest condition while walking in the exoskeleton without assistance. This may have added to errors in metabolic estimates of the unassisted walking condition at 15 degrees.

#### 2) Torques and Kinematics

Applied exoskeleton torques were measured using strain gauges at the ankles and were calculated using load cells and known lever arms for the hips and knees [42]. Exoskeleton joint angles were measured to estimate user kinematics, assuming only small amounts of relative movement between the user and the exoskeleton. Stride frequency was calculated using vertical ground reaction forces measured by the instrumented treadmill (Bertec). Measurements were averaged across both legs.

#### 3) Exoskeleton power

Applied exoskeleton power was calculated by multiplying the measured torque by joint velocity. Joint velocities were calculated by differentiating measured joint angles and low-pass filtering at 50 Hz. Average power was calculated for each joint. Positive power and negative power were calculated by taking the average of the positive power values and the negative power values, respectively. Each joint power is the sum of both legs, and total power was the sum of all joint powers.

#### 4) Muscle Activity

We used surface EMG (Delsys Trigno) to measure muscle activity. We measured the soleus, gastrocnemius lateralis, vastus lateralis, and gluteus maximus of the participant’s right leg (which was the dominant leg of 2 participants). Signals were passed through a 3rd order bandpass filter of 40 to 450 Hz, rectified, and then smoothed with a 3rd order low pass filter of 10 Hz [48]. We subtracted the baseline noise offset, then normalized the activity of each muscle in each condition to the maximum of the walking in the exoskeleton without torque at that incline. For example, soleus activity for walking with assistance at 5 degrees was normalized to the peak activity of walking in the exoskeleton without assistance at 5 degrees. This was done to identify the effect of assistance, and because each incline level was validated on a separate day. The sensor locations are similar to the protocol of previous gait analysis experiments [49] with slight adjustments to avoid interference from straps and device contacts.

## III. Results

### A. Metabolic Cost and Cost of Transport

The cost of transport and the metabolic cost of walking were reduced in all assisted conditions (Fig. 2). To evaluate the metabolic cost of walking, the metabolic cost of quiet standing for each participant was subtracted from the measured metabolic costs of participant’s walking conditions. For walking without the exoskeleton, metabolic cost increased from level-ground walking to 10 degrees and was similar between 10 and 15 degrees, because increases in incline level and decreases in walking speed both had an effect on metabolic cost (A-C) [16], [17], [18], [50]. To evaluate the effect of incline specifically, we calculated and compared the cost of transport by dividing the metabolic cost of walking by the walking speed. Cost of transport for walking without the exoskeleton increased linearly as incline level increased (R^2^ = 0.94, A-D).

**Fig. 2.**
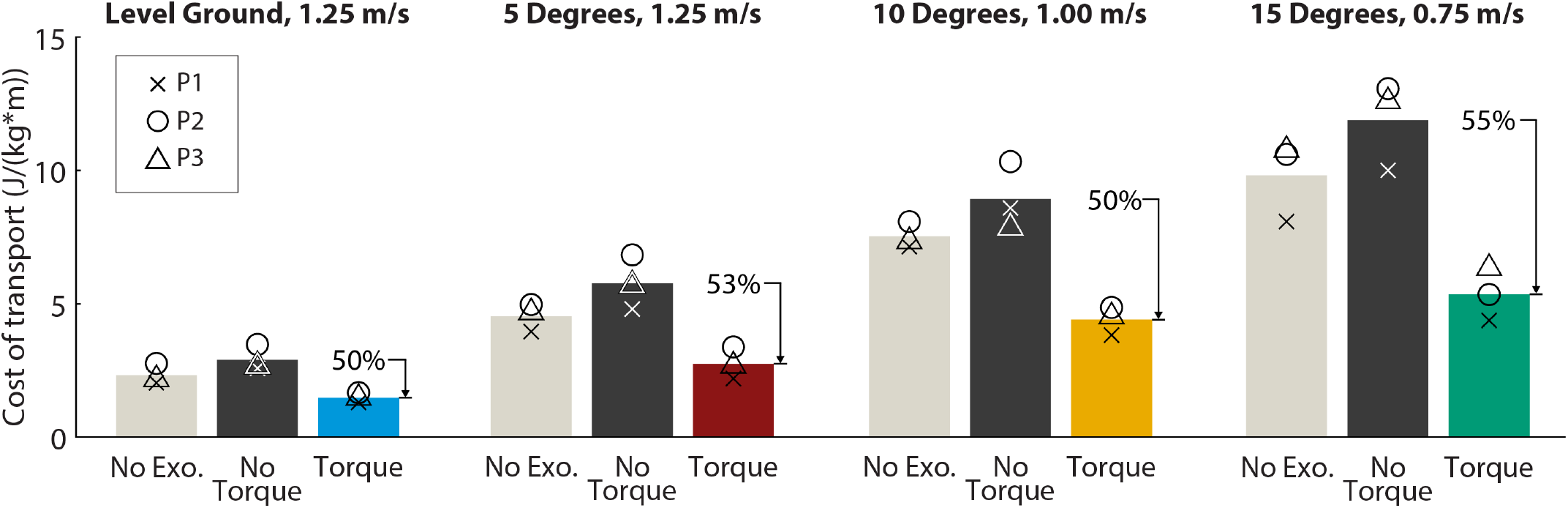
Cost of transport. Cost of transport (J / (kg*m)) for each condition (no exoskeleton: gray, no torque: black, optimized assistance: color). Normalizing by speed shows an increase in cost of transport as incline increases. Percent reduction relative to walking in the exoskeleton without assistance is shown above each optimized assistance bar. Individual participant values are indicated by the symbols.

For all assistance conditions, cost of transport was reduced by at least 50% on average relative to walking in the device with no assistance (Fig. 2). Exoskeleton assistance reduced the cost of transport relative to unassisted walking by 50% for the level-ground condition (range 46%-53%, p = 0.02), by 53% for the 5-degree condition (range 51%-54%, p = 0.01), by 50% for the 10-degree condition (range 42%-56%, p = 0.02), and by 55% for the 15-degree condition (range 50%-59%, p = 0.01). This corresponds to large absolute reductions in metabolic cost, with the cost of walking on 15 degrees reduced by 4.9 W/kg (range: 4.2 to 5.8 W/kg, A-C).

Relative to walking without the exoskeleton, this corresponds to a reduction in cost of transport of 37% for the level-ground condition (range 34%-41%, p = 0.02), by 40% for the 5-degree condition (range 32%-44%, p = 0.004), by 42% for the 10-degree condition (range 38%-47%, p = 0.002), and by 46% for the 15-degree condition (range 41%-50%, p = 0.01). This comparison to walking with no exoskeleton does not include the full effects of added mass of a mobile device, which might be larger than our emulator due to the need to carry motors and batteries. Nevertheless, these large reductions indicate that whole-leg exoskeleton assistance can greatly reduce the energy the user consumes during incline walking across a range of grades, even as walking speed decreases.

### B. Exoskeleton Torque

Exoskeleton extension torque magnitudes tended to increase with steeper inclines, often increasing until hitting comfort-based limits at 10 and 15 degrees (Fig. 3, A-E). For all joints, timing parameters were often consistent across participants and grades. For example, the timing of peak hip extension torque ranged from 9.1% to 11.5% of stride (measured from heel strike) across all participants and conditions. For torque magnitudes, peak hip extension torque magnitudes tended to increase with incline, often until reaching comfort-based limits at 10 and 15 degrees (0.9 Nm/kg). Knee extension torque tended to similarly increase. For two participants, knee stiffness increased with incline until reaching maximum allowed levels, while one participant had consistent optimized knee stiffness across grades. While all participants optimized to larger knee extension torques at an incline than over level-ground, there was inter-participant variability in optimal knee extension magnitudes, with peak torques ranging from 0.35 Nm/kg to 0.7 Nm/kg for walking at 10 degrees (A-E). Knee flexion torque during damping control in late swing decreased for steeper inclines, possibly due to decreases in walking speed and stride frequency. Ankle torque magnitude often reached comfort-based limits, with two participants optimizing to maximum-allowed torque for all conditions. For incline walking, we relaxed the ankle torque constraint from 0.8 Nm/kg to 0.9 Nm/kg, as allowed by participants’ comfort. For one participant, ankle torque optimized to a smaller value at 15 degrees compared to other conditions. While subject-specific differences in magnitudes were present, hip and knee extension torques generally increased with incline as expected based on biological torques [31], while large ankle plantarflexion torques at comfort-based limits were effective across conditions.

**Fig. 3.**
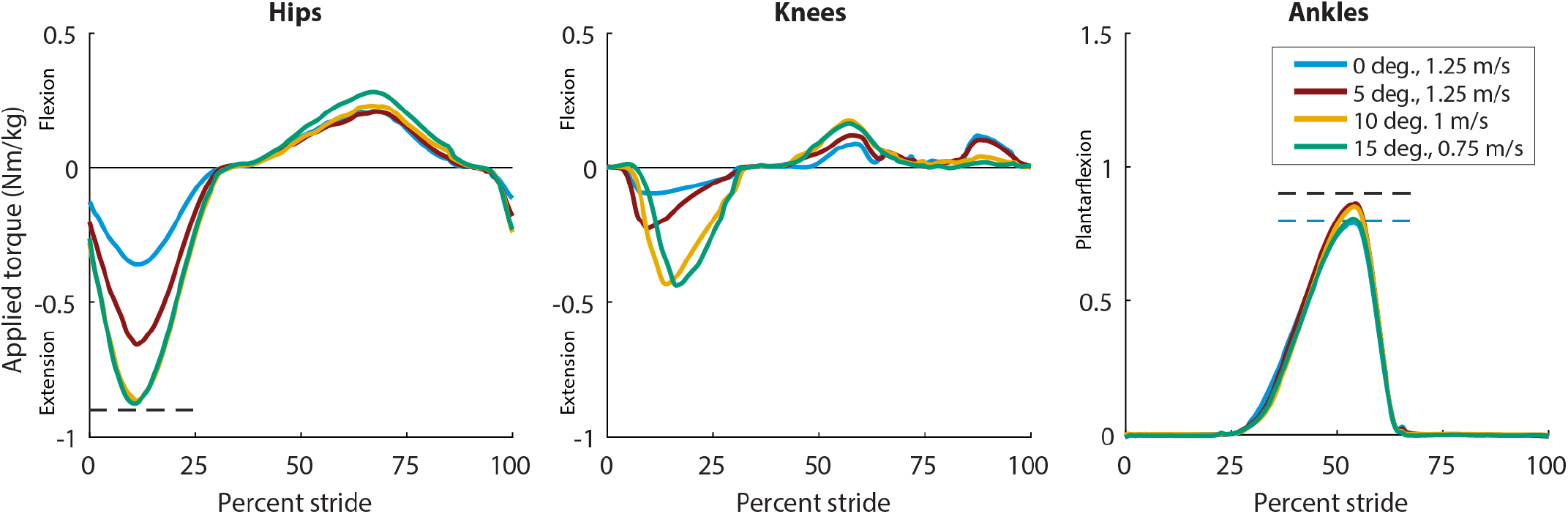
Applied Exoskeleton Torques. Optimized assistance applied to the exoskeleton for each condition. Dashed lines for hip extension and ankle plantarflexion indicate maximum allowed torques that were approached during the optimization. For the ankles, blue dashed line indicates max torque for level-ground walking (0.8 Nm/kg), which was relaxed for incline (black dashed line, 0.9 Nm/kg) because changes in kinematics allowed larger torques to still be comfortable. Knee extension assistance was restricted based on virtual spring stiffness.

### C. Exoskeleton Power

Total exoskeleton power tended to increase with incline, with shifting contributions from each joint (Fig. 4, A-F). Total exoskeleton power and total positive exoskeleton power increased with incline up to 10 degrees (level vs 5 degrees: p = 0.02, 5 vs 10 degrees: p = 0.04) with similar powers between 10 and 15 degrees (p = 0.74). For the hips and knees, positive power increased as incline increased, although changes were not always statistically significant (A-F). For the ankles, positive power was largest at 5 degrees, and decreased as incline increased to 10 and 15 degrees. Powers varied across participants at the joint level, reflecting differences in optimized torque values or potentially indicating different walking strategies between participants. For example, participant 3 optimized to smaller ankle torques and larger knee torques than the other participants. Overall, total positive exoskeleton power in W/kg trended similarly to reduction in metabolic cost in W/kg (Fig. 4, right), indicating a potential relationship between power delivered and metabolic reduction.

**Fig. 4.**
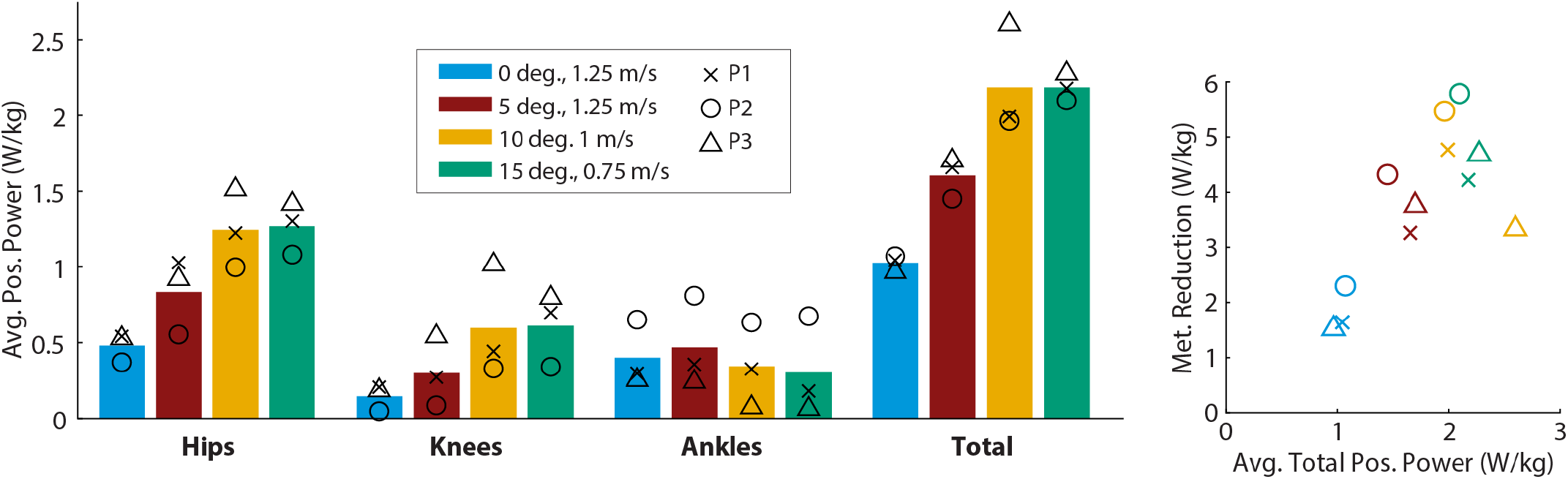
Applied positive exoskeleton powers. Exoskeleton applied positive power at each joint and total positive power for each condition. Positive power was calculated as the average of all powers greater than zero during the average stride. (Right) Absolute metabolic reduction (W/kg) plotted as a function of total positive power (W/kg). Individual participant values are indicated by the symbols.

### D. Kinematics

With assistance, users tended to walk with more extended joints, usually consistent with directions of applied torque (Fig. 5, A-G). Unassisted kinematics changed with incline level as expected based on previous studies [51]: as incline increased, users walked with more hip flexion, knee flexion, and ankle dorsiflexion. When assistance torques were applied, users’ kinematics changed, showing more rapid extension of the hips and knees during stance. For the hips, assistance increased the rate of hip extension in early stance, consistent with the direction and timing of applied torque. While the rate of extension changed, maximum hip extension angle did not consistently change with assistance. For hip flexion, maximum flexion angle in swing (70% to 100% stride) was decreased with assistance for incline conditions, even though the device actively assisted hip flexion, possibly due to changes in stride frequency. For the knees, assistance increased the rate of knee extension in early stance, consistent with the direction of applied torque. Around the timing of push-off and during swing, assistance increased knee flexion angle. For the ankles, there was slightly less dorsiflexion during stance with assistance, but there were no large changes to maximum plantarflexion angle despite the presence of large ankle torques.

**Fig. 5.**
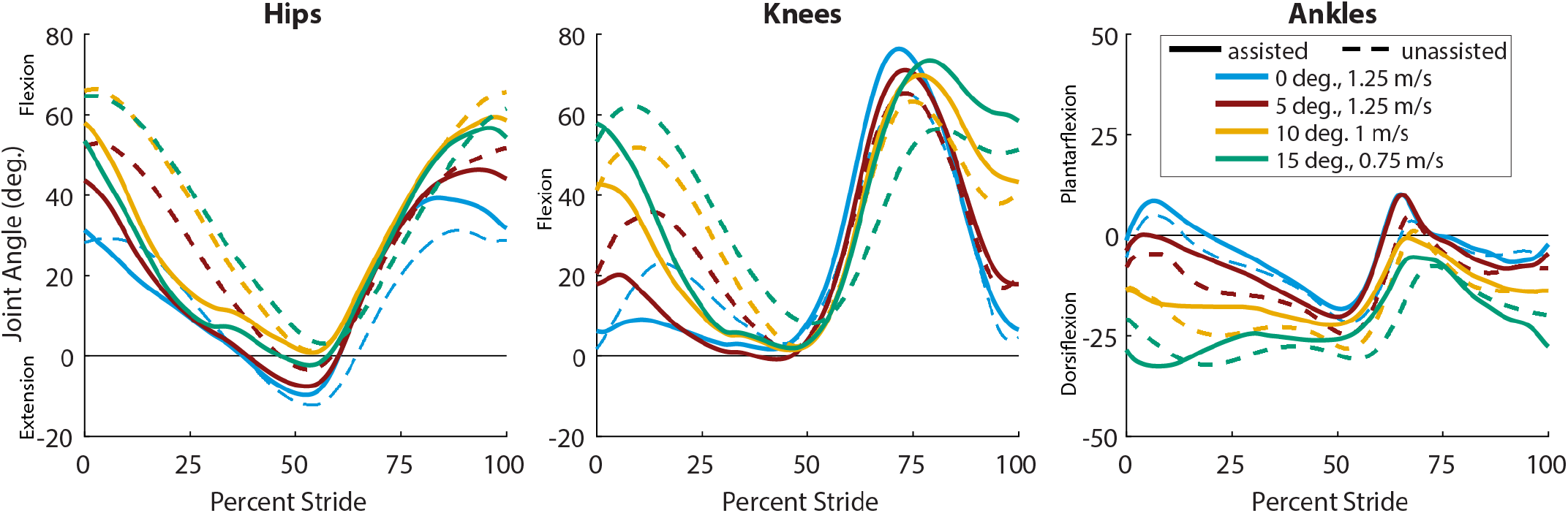
Average joint kinematics. Average joint angle as a percentage of stride at the hips (left), knees (center), and ankles (right) for each condition. Shown here are the average for both legs across all participants (N = 3). Unassisted walking for each condition is shown with dashed lines, assisted walking is shown with solid lines. Extension assistance at the hips and knees demonstrate increased extension rate during stance.

Both incline level and exoskeleton assistance seemed to affect stride frequency, but changes were not statistically significant (A-H). For unassisted walking, stride frequency decreased as speed decreased and incline increased. Exoskeleton assistance increased stride frequency, with small changes at level-ground walking, and larger increases for steeper inclines. While trends were consistent, there were large inter-participant differences in stride frequencies, likely leading to statistical insignificance. For example, for unassisted walking at 15 degrees, stride frequencies ranged from 0.52 Hz to 0.75 Hz across participants.

### E. Muscle Activity

Exoskeleton assistance reduced muscle activity relative to unassisted walking for a number of muscles across conditions and participants (Fig. 6, A-I). Each incline condition was tested on a separate day. For each muscle at each incline level, activity was normalized to the peak activity of that muscle during the unassisted condition to evaluate the effect of assistance. For the gluteus maximus, assistance reduced activity during stance when hip extension assistance was applied. For the vastus lateralis, activity was reduced during knee extension assistance in stance, with larger reductions in activity for incline conditions than for level-ground walking. During level-ground walking, assistance increased vastus lateralis activity around the time of toe-off, but this increase in activity was not seen for the incline conditions. For the soleus and gastrocnemius, activity was reduced by exoskeleton assistance around push-off. This was consistent across participants and coincides with the timing of ankle plantarflexion assistance and knee flexion assistance, but could also be related to hip extension assistance [38]. These changes in muscle activity indicate assistance can be effective across incline levels.

**Fig. 6.**
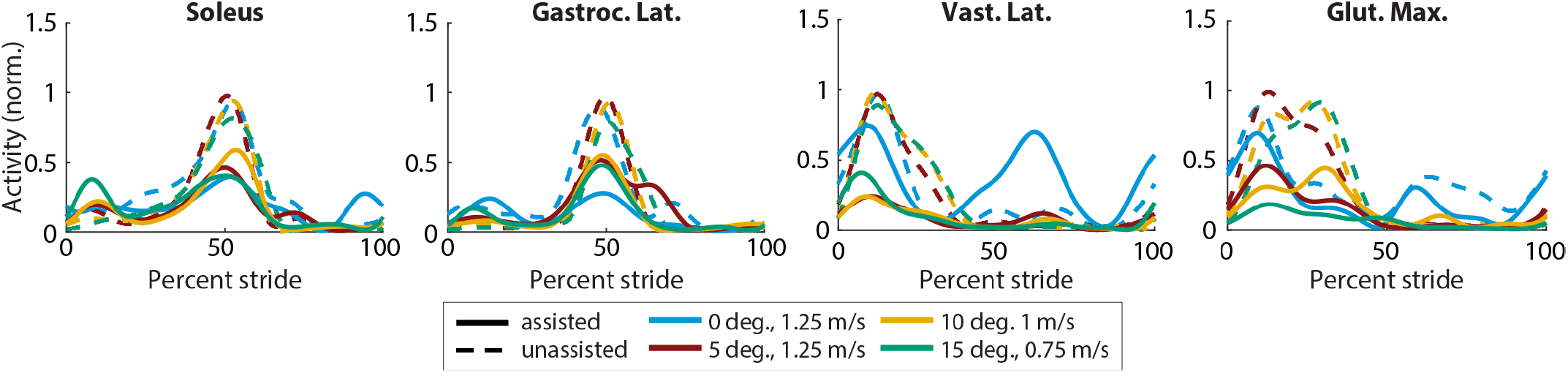
Average muscle activity. Muscle activity measured during walking using surface EMG for each condition. Lines shown are the average across all participants (N = 3). The EMG signal was filtered, averaged across all strides, had baseline activity removed to eliminate noise, and normalized to the peak value of walking in the exoskeleton without assistance for the given incline and speed (dashed).

## IV. Discussion

Whole-leg exoskeleton assistance can deliver large improvements to walking across a range of inclines. Assistance reduced metabolic cost by at least 50% relative to walking without assistance at every incline level, indicating that devices could be successful in a wide range of terrain and environments. In terms of absolute reductions in metabolic cost, the most strenuous condition was reduced by around 4.9 W/kg, more than twice the entire cost of level walking. Comparing absolute reductions in cost, the previous best improvement to incline walking was a reduction of 0.73 W/kg at 5 degrees from hip-only assistance by an untethered device [29]. These results indicate that, for expert users with significant device capabilities, exoskeletons could greatly improve walking performance on inclines.

The magnitude of these metabolic cost reductions may help foster device adoption. The demonstrated metabolic reductions were larger than the just-noticeable difference of metabolic cost of 20% [52], indicating users could perceive the benefit of assistance. For the most strenuous conditions, unassisted walking was difficult and sometimes verged into the anaerobic region of activity (respiratory quotient greater than 1). Large improvements from assistance kept users in the aerobic region, which would likely improve user endurance. These large benefits indicate that assisting inclines or other strenuous tasks may deliver noticeable benefits to walking for able-bodied users who may not perceive the need for assistance on level-ground.

Optimized exoskeleton assistance applied more positive power as incline increased, corresponding to larger absolute metabolic reductions. Total exoskeleton power increased as incline increased up to 10 degrees, consistent with changes in biological joint work and power with incline in unassisted walking [30], [31]. To compare to biological values, total positive exoskeleton power at 5 degrees (1.60 W/kg) was slightly less than total positive biological power in unassisted walking at 5.71 degrees (1.71 W/kg) [31]. The distribution by joint was similar, with powers of 0.83, 0.30, and 0.47 W/kg for the exoskeleton hip, knee, and ankle, compared to biological powers of 0.81, 0.32, and 0.58 W/kg for the same joints during walking at the same speed on an incline of 5.71 degrees [31]. Trends in exoskeleton power indicate that more power is optimal for steeper inclines and corresponds to larger absolute reductions in metabolic cost. However, this does not necessarily mean that any particular assistance condition would have benefited from more applied positive power [53]; if it had, the optimizer could have increased torque magnitudes if not yet at comfort-based limits (e.g. hip extension torque during level-ground walking could have increased). Alternatively, a user could also have chosen a coordination pattern with more joint range of motion during exoskeleton torque application. Instead, it implies that larger exoskeleton powers are more optimal in scenarios where costs are higher and where biological powers are higher, indicating potential for larger assistance.

Trends in exoskeleton power with incline level were not consistent across joints. Based on biological data, we expected increases in exoskeleton positive power at all three joints as incline increased [30], [31]. Exoskeleton positive power increased for the hips and knees with incline, as expected. Ankle positive power decreased when walking at 10 and 15 degrees. Based on biological trends, we expected the ankle’s contribution to total power to shrink [31], but we did not expect a decrease in power. This decrease was most prominent for one participant (P3), who converged to smaller ankle torque magnitudes at 10 and 15 degrees. This participant and one other participant (P2) reported some discomfort at the ankles during assistance, and decreases in power may be related to kinematic adaptations to reduce discomfort at the joint. These decreases may also be related to decreases in walking speed, because we expect smaller powers at slower speeds.

These results suggest ways of designing better exoskeleton controllers and hardware for varying terrain. Our results indicate that as incline increases, devices should deliver more torque in hip extension and knee extension, while consistently applying large ankle plantarflexion torques. Exoskeleton controllers could apply continuous control with changing incline by interpolating device parameters between the discretely optimized incline levels. Considering timing parameters were consistent across users and inclines, devices could apply a static timing profile while optimizing torque magnitudes in real-time. Future work could use exoskeleton sensor data to detect incline level [54] and update their control in real time to apply optimal assistance over varying inclines in unstructured terrain. The optimized exoskeleton peak torque magnitudes were larger than the capabilities of many existing mobile devices [42]. Exoskeleton designers who want to maximally assist walking could use these optimized magnitudes to inform the design of future devices with larger torque and power capabilities to optimally assist walking on steep inclines. Assisting the whole-leg produced larger reductions than previous single-joint attempts at assisting incline walking, consistent with our direct comparisons on level-ground [38]. This indicates that exoskeleton designers who want to deliver maximum reductions to users could consider assisting the entire leg. With these design specifications and future improvements to control strategies, mobile devices are poised to be successful in the field.

The results of this study could help us understand how users adapt to and benefit from exoskeletons, potentially improving our models of motor control. Changes in kinematics and stride frequency indicate that users could be adapting their walking strategy to best take advantage of the assistance. For example, users extended their hips and knees earlier in stance with assistance, possibly to raise their center of gravity earlier in stride to reduce activity of knee extensors [28]. Understanding how users adapt their kinematics in the presence of exoskeleton torques could improve our neuromuscular models of walking control and inform how musculoskeletal simulations of assistance could consider kinematic changes [55].

This study increases the range of exoskeleton powers and metabolic cost reductions delivered by exoskeletons, which could help us understand how users benefit from assistance. The larger absolute reductions in this study relative to level-ground assistance could be due to the intrinsic energy requirements of the task of walking uphill. More work is done by the muscles against gravity as incline increases, giving more opportunity for the device to assist, whereas there are no intrinsic work requirements for level-ground walking. While absolute reductions changed, future work could try to understand why percent reductions were similar across conditions. For level-ground walking, we previously hypothesized that some of the remaining energy was necessary for balance or could be related to the lack of assistance in the frontal plane. Because metabolic reductions were close to 50% for incline conditions as well, this could indicate that the cost to balance in the frontal plane increases as either incline increases, speed decreases, or as applied torque magnitudes increase. Muscle activity reductions were similar across conditions, indicating there may be some percentage of muscle activity needed for the user to share a meaningful amount of control, or that some activity must remain to enable fast reactions for balance [7]. It could be that we could deliver larger percent reductions if torque magnitudes were not limited.

Our study was limited by the number of participants and related effects of the COVID-19 pandemic, but we expect the large improvements demonstrated to extend to more users. This was a long study conducted in part with other optimization experiments [37], [38] during difficult external conditions (the COVID-19 pandemic). As such, we were only able to complete three participants for the study. However, given the magnitude of the metabolic changes and the consistency of the responses across participants, this sample size is sufficient to identify that whole-leg assistance delivers significant improvements to metabolic cost across incline levels. Given three participants and a desired statistical power of 0.8, and assuming metabolic reductions have a standard deviation of 7.3% [7], we can confidently detect metabolic reductions of 24% and larger [38]. The COVID-19 pandemic also meant participants wore cloth masks underneath their metabolic masks, which decreased the accuracy of the metabolic estimates. We expected a metabolic cost of walking of 6.56 W/kg for walking at 5 degrees ([16] interpolating from 7.03 W/kg at 5.71 degrees), whereas we measured a cost of 5.66 W/kg for walking without the exoskeleton at 5 degrees. While the absolute measurements may be affected, we expect that the percent reductions should be accurate since the mask is affecting all incline conditions [37], [38].

This study could have been improved by longer optimizations, testing additional controller parameterizations, or by testing self-selected walking speed. Although we gave our users substantial optimization time for each incline level, more time may have improved the outcomes by improving likelihood of optimizer convergence, as well as by giving the user more training for each condition. For many of the walking conditions, torques optimized to the allowed limits based on comfort for all three joints. Improving our control strategy to reduce user discomfort and allow larger torques could lead to larger reductions in metabolic cost. This would also allow a better trend to be seen between optimal assistance and incline level. By varying speed and incline, we were able to maximize functional relevance and also ensure users were in the aerobic region while walking to ensure accurate metabolic measurements. However, if we were able to hold walking speed constant, or allow users to walk at a self-selected speed, we may better be able to identify the effect of incline specifically. Lastly, we validated each of the conditions separately after each optimization. While this is best practice in terms of consolidating learning and mitigating effects of fatigue, we were unable to record muscle activations across all inclines on the same day, and variability between EMG attachment between days means we are unable to directly compare how muscle activity changed as incline changed.

These results suggest new opportunities for future study and development, including new control strategies, cost functions, user populations, and mobile devices. Having identified optimal torques and powers for walking on inclines, future studies could see how these results translate to mobile devices and could use those systems to study incline assistance in unstructured environments where inclines are common, such as in hilly or rocky terrain. Future work could study how assistance could increase walking speeds or load carriage capabilities while walking on inclines. Often, torques converged on the limits of what we could comfortably apply. More investigation into interface design and a better biomechanical understanding of joint discomfort could lead to devices applying greater assistance levels comfortably, which could result in greater reductions. While the large optimized assistance magnitudes seem consistent with changes in biological torques and powers with incline, it may be possible to get similar reductions with smaller torques or powers. Future work could optimize for reducing metabolic cost while also penalizing large actuation requirements, which are costly for mobile devices. By reducing energy costs at large inclines, assistance could increase the self-selected speed of people walking on steep terrain. Future work could also allow for walking speed to vary with incline using a self-paced treadmill [56], and could optimize for some combination of speed and metabolic cost. While our users did not need balance assistance in this study, additional studies could investigate how to assist balance, especially over uneven, inclined terrain. Finally, while our findings demonstrate that exoskeleton assistance is effective for young, able-bodied users, an important continuation of this research is to extend assistance to older adults and people with disabilities. Continuing to emphasize the ability to traverse challenging environmental elements, such as ramps and stairs, will play a significant role in bringing exoskeleton use into the daily life of those who would benefit from it.

## Appendix A

The appendix information are available as a separate supplementary document. The contents of this supplementary are listed below.

A. *Parameterization of Torque Control*
B. *Effect of Cloth Mask on Metabolic Cost Measurements*
C. *Metabolic Cost for Each Condition*
D. *Cost of Transport Table*
E. *Optimized Torques for Each Participant*
F. *F. Applied Exoskeleton Powers*
G. *Kinematics for Each Participant*
H. *Stride Frequency*
I. *Muscle Activity for Each Participant*

## Acknowledgment

The authors would like to thank the DEVCOM Soldier Center for support of this work, N. Bianco for assistance in controller development, R. Martin and A. Lakmazaheri for feedback on this manuscript, and all of the Stanford Biomechatronics Lab for their feedback and support. We would also like to thank the staff and administrators who were able to reopen the lab during the COVID-19 pandemic.

## References

[1] M. Y. Zarrugh, F. N. Todd, and H. J. Ralston, “Optimization of energy expenditure during level walking,” European Journal of Applied Physiology and Occupational Physiology, vol. 33, no. 4, pp. 293–306, Dec. 1974. [Online]. Available: https://doi.org/10.1007/BF00430237

[2] G. S. Sawicki, O. N. Beck, I. Kang, and A. J. Young, “The exoskeleton expansion: improving walking and running economy,” Journal of NeuroEngineering and Rehabilitation, vol. 17, no. 1, p. 25, Feb. 2020. [Online]. Available: https://doi.org/10.1186/s12984-020-00663-9

[3] P. Malcolm, W. Derave, S. Galle, and D. D. Clercq, “A Simple Exoskeleton That Assists Plantarflexion Can Reduce the Metabolic Cost of Human Walking,” PLOS ONE, vol. 8, no. 2, p. e56137, Feb. 2013. [Online]. Available: https://journals.plos.org/plosone/article?id=10.1371/journal.pone.0056137

[4] L. M. Mooney, E. J. Rouse, and H. M. Herr, “Autonomous exoskeleton reduces metabolic cost of human walking during load carriage,” Journal of NeuroEngineering and Rehabilitation, vol. 11, no. 1, p. 80, May 2014. [Online]. Available: https://doi.org/10.1186/1743-0003-11-80

[5] S. H. Collins, M. B. Wiggin, and G. S. Sawicki, “Reducing the energy cost of human walking using an unpowered exoskeleton,” Nature, vol. 522, no. 7555, pp. 212–215, Jun. 2015. [Online]. Available: http://www.nature.com/articles/nature14288

[6] K. Seo, J. Lee, Y. Lee, T. Ha, and Y. Shim, “Fully autonomous hip exoskeleton saves metabolic cost of walking,” in 2016 IEEE International Conference on Robotics and Automation (ICRA),May 2016, pp. 4628–4635.

[7] J. Zhang, P. Fiers, K. A. Witte, R. W. Jackson, K. L. Poggensee, C. G. Atkeson, and S. H. Collins, “Human-in-the-loop optimization of exoskeleton assistance during walking,” Science, vol. 356, no. 6344, pp. 1280–1284, Jun. 2017, publisher: American Association for the Advancement of Science Section: Report. [Online]. Available: http://science.sciencemag.org/content/356/6344/1280

[8] B. T. Quinlivan, S. Lee, P. Malcolm, D. M. Rossi, M. Grimmer, C. Siviy, N. Karavas, D. Wagner, A. Asbeck, I. Galiana, and C. J. Walsh, “Assistance magnitude versus metabolic cost reductions for a tethered multiarticular soft exosuit,” Science Robotics, vol. 2, no. 2, Jan. 2017, publisher: Science Robotics Section: Research Article. [Online]. Available: http://robotics.sciencemag.org/content/2/2/eaah4416

[9] Y. Ding, M. Kim, S. Kuindersma, and C. J. Walsh, “Human-in-the-loop optimization of hip assistance with a soft exosuit during walking,” Science Robotics, vol. 3, no. 15, p. eaar5438, Feb. 2018. [Online]. Available: https://robotics.sciencemag.org/content/3/15/eaar5438

[10] S. Lee, J. Kim, L. Baker, A. Long, N. Karavas, N. Menard Galiana, and C. J. Walsh, “Autonomous multi-joint soft exosuit with augmentation-power-based control parameter tuning reduces energy cost of loaded walking,” Journal of NeuroEngineering and Rehabilitation, vol. 15, no. 1, p. 66, Dec. 2018. [Online]. Available: https://jneuroengrehab.biomedcentral.com/articles/10.1186/s12984-018-0410-y

[11] P. Malcolm, S. Galle, W. Derave, and D. De Clercq, “Bi-articular Knee-Ankle-Foot Exoskeleton Produces Higher Metabolic Cost Reduction than Weight-Matched Mono-articular Exoskeleton,” Frontiers in Neuroscience, vol. 12, 2018, publisher: Frontiers. [Online]. Available: https://www.frontiersin.org/articles/10.3389/fnins.2018.00069/full

[12] B. Lim, J. Lee, J. Jang, K. Kim, Y. J. Park, K. Seo, and Y. Shim, “Delayed Output Feedback Control for Gait Assistance With a Robotic Hip Exoskeleton,” IEEE Transactions on Robotics, vol. 35, no. 4, pp. 1055–1062, Aug. 2019. [Online]. Available: https://doi.org/10.1109/TRO.2019.2913318

[13] M. K. MacLean and D. P. Ferris, “Energetics of Walking With a Robotic Knee Exoskeleton,” Journal of Applied Biomechanics, vol. 35, no. 5, pp. 320–326, Oct. 2019.

[14] W. Cao, C. Chen, H. Hu, K. Fang, and X. Wu, “Effect of Hip Assistance Modes on Metabolic Cost of Walking With a Soft Exoskeleton,” IEEE Transactions on Automation Science and Engineering, pp. 1–11, 2020, conference Name: IEEE Transactions on Automation Science and Engineering.

[15] J. Young, J. Foss, H. Gannon, and D. P. Ferris, “Influence of Power Delivery Timing on the Energetics and Biomechanics of Humans Wearing a Hip Exoskeleton,” Frontiers in Bioengineering and Biotechnology, vol. 5, Mar. 2017. [Online]. Available: https://www.ncbi.nlm.nih.gov/pmc/articles/PMC5340778/

[16] Silder, T. Besier, and S. L. Delp, “Predicting the Metabolic Cost of Incline Walking from Muscle Activity and Walking Mechanics,” Journal of biomechanics, vol. 45, no. 10, pp. 1842–1849, Jun. 2012. [Online]. Available: https://www.ncbi.nlm.nih.gov/pmc/articles/PMC4504736/

[17] E. Minetti, C. Moia, G. S. Roi, D. Susta, and G. Ferretti, “Energy cost of walking and running at extreme uphill and downhill slopes,” Journal of Applied Physiology, vol. 93, no. 3, pp. 1039–1046, Sep. 2002, publisher: American Physiological Society. [Online]. Available: https://journals.physiology.org/doi/full/10.1152/japplphysiol.01177.2001

[18] N. A. Pimental and K. B. Pandolf, “Energy expenditure while standing or walking slowly uphill or downhill with loads,” Ergonomics, vol. 22, no. 8, pp. 963–973, Aug. 1979, publisher: Taylor & Francis eprint: https://doi.org/10.1080/00140137908924670. [Online]. Available: https://doi.org/10.1080/00140137908924670

[19] R. M. Orr, R. Pope, V. Johnston, and J. Coyle, “Soldier occupational load carriage: a narrative review of associated injuries,” International Journal of Injury Control and Safety Promotion, vol. 21, no. 4, pp. 388–396, Oct. 2014, publisher: Taylor & Francis eprint: https://doi.org/10.1080/17457300.2013.833944. [Online]. Available: https://doi.org/10.1080/17457300.2013.833944

[20] S. Hignett, J. W. Willmott, and S. Clemes, “Mountain rescue stretchers: Usability trial,” Work, vol. 34, no. 2, pp. 215–222, Jan. 2009, publisher: IOS Press. [Online]. Available: https://content.iospress.com/articles/work/wor00918

[21] P. E. Martin, D. E. Rothstein, and D. D. Larish, “Effects of age and physical activity status on the speed-aerobic demand relationship of walking,” Journal of Applied Physiology (Bethesda, Md.: 1985), vol. 73, no. 1, pp. 200–206, Jul. 1992.

[22] O. S. Mian, J. M. Thom, L. P. Ardigò, M. V. Narici, and A. E. Minetti, “Metabolic cost, mechanical work, and efficiency during walking in young and older men,” Acta Physiologica (Oxford, England), vol. 186, no. 2, pp. 127–139, Feb. 2006.

[23] F. Lauretani, C. R. Russo, S. Bandinelli, B. Bartali, C. Cavazzini, A. Di Iorio, A. M. Corsi, T. Rantanen, J. M. Guralnik, and L. Ferrucci, “Age-associated changes in skeletal muscles and their effect on mobility: an operational diagnosis of sarcopenia,” Journal of Applied Physiology (Bethesda, Md.: 1985), vol. 95, no. 5, pp. 1851–1860, Nov. 2003.

[24] S. Song and H. Geyer, “Predictive neuromechanical simulations indicate why walking performance declines with ageing,” The Journal of Physiology, vol. 596, no. 7, pp. 1199–1210, 2018, eprint: https://physoc.onlinelibrary.wiley.com/doi/pdf/10.1113/JP275166. [Online]. Available: https://physoc.onlinelibrary.wiley.com/doi/abs/10.1113/JP275166

[25] J. R. Franz and R. Kram, “Advanced age and the mechanics of uphill walking: A joint-level, inverse dynamic analysis,” Gait & Posture, vol. 39, no. 1, pp. 135–140, Jan. 2014. [Online]. Available: https://www.sciencedirect.com/science/article/pii/S0966636213002932

[26] J. D. Ortega and C. T. Farley, “Effects of aging on mechanical efficiency and muscle activation during level and uphill walking,” Journal of Electromyography and Kinesiology, vol. 25, no. 1, pp. 193–198, Feb. 2015. [Online]. Available: https://www.sciencedirect.com/science/article/pii/S1050641114001898

[27] G. S. Sawicki and D. P. Ferris, “Mechanics and energetics of incline walking with robotic ankle exoskeletons,” Journal of Experimental Biology, vol. 212, no. 1, pp. 32–41, Jan. 2009, publisher: The Company of Biologists Ltd Section: Research Article. [Online]. Available: https://jeb.biologists.org/content/212/1/32

[28] S. Galle, P. Malcolm, W. Derave, and D. De Clercq, “Uphill walking with a simple exoskeleton: Plantarflexion assistance leads to proximal adaptations,” Gait & Posture, vol. 41, no. 1, pp. 246–251, Jan. 2015. [Online]. Available: https://www.sciencedirect.com/science/article/pii/S0966636214007413

[29] K. Seo, J. Lee, and Y. J. Park, “Autonomous hip exoskeleton saves metabolic cost of walking uphill,” in 2017 International Conference on Rehabilitation Robotics (ICORR), Jul. 2017, pp. 246–251, iSSN: 1945-7901.

[30] J. R. Montgomery and A. M. Grabowski, “The contributions of ankle, knee and hip joint work to individual leg work change during uphill and downhill walking over a range of speeds,” Royal Society Open Science, vol. 5, no. 8, p. 180550, Aug. 2018. [Online]. Available: https://royalsocietypublishing.org/doi/10.1098/rsos.180550

[31] R. W. Nuckols, K. Z. Takahashi, D. J. Farris, S. Mizrachi, R. Riemer, and G. S. Sawicki, “Mechanics of walking and running up and downhill: A joint-level perspective to guide design of lower-limb exoskeletons,” PLOS ONE, vol. 15, no. 8, p. e0231996, Aug. 2020, publisher: Public Library of Science. [Online]. Available: https://journals.plos.org/plosone/article?id=10.1371/journal.pone.0231996

[32] J. R. Franz, N. E. Lyddon, and R. Kram, “Mechanical work performed by the individual legs during uphill and downhill walking,” Journal of Biomechanics, vol. 45, no. 2, pp. 257–262, Jan. 2012. [Online]. Available: https://linkinghub.elsevier.com/retrieve/pii/S0021929011006762

[33] J. C. Wall, J. W. Nottrodt, and J. Charteris, “The effects of uphill and downhill walking on pelvic oscillations in the transverse plane,” Ergonomics, vol. 24, no. 10, pp. 807–816, Oct. 1981, publisher: Taylor & Francis eprint: https://doi.org/10.1080/00140138108924901. [Online]. Available: https://doi.org/10.1080/00140138108924901

[34] K. Kawamura, A. Tokuhiro, and H. Takechi, “Gait analysis of slope walking: a study on step length, stride width, time factors and deviation in the center of pressure.” Acta Medica Okayama, vol. 45, no. 3, pp. 179–184, Jun. 1991, publisher: Okayama University Medical School. [Online]. Available: http://ousar.lib.okayama-u.ac.jp/32212

[35] J. Sun, M. Walters, N. Svensson, and D. Lloyd, “The influence of surface slope on human gait characteristics: a study of urban pedestrians walking on an inclined surface,” Ergonomics, vol. 39, no. 4, pp. 677–692, Mar. 1996. [Online]. Available: http://www.tandfonline.com/doi/abs/10.1080/00140139608964489

[36] R. Castano and H. J. Huang, “Speed-related but not detrended gait variability increases with more sensitive self-paced treadmill controllers at multiple slopes,” PLOS ONE, vol. 16, no. 5, p. e0251229, May 2021, publisher: Public Library of Science. [Online]. Available: https://journals.plos.org/plosone/article?id=10.1371/journal.pone.0251229

[37] G. M. Bryan, P. W. Franks, S. Song, A. S. Voloshina, Reyes, M. P. O’Donovan, K. N. Gregorczyk, and S. H. Collins, “Optimized hip-knee-ankle exoskeleton assistance at a range of walking speeds,” bioRxiv, Mar. 2021. [Online]. Available: http://biorxiv.org/lookup/doi/10.1101/2021.03.26.437212

[38] P. W. Franks, G. M. Bryan, R. M. Martin, R. Reyes, and H. Collins, “Comparing optimized exoskeleton assistance of the hip, knee, and ankle in single and multi-joint configurations,” bioRxiv, p. 2021.02.19.431882, Feb. 2021, publisher: Cold Spring Harbor Laboratory Section: New Results. [Online]. Available: https://www.biorxiv.org/content/10.1101/2021.02.19.431882v1

[39] Lee, B. J. McLain, I. Kang, and A. J. Young, “Biomechanical Comparison of Assistance Strategies Using a Bilateral Robotic Knee Exoskeleton,” IEEE Transactions on Biomedical Engineering, vol. 68, no. 9, pp. 2870–2879, Sep. 2021.

[40] L. Dembia, A. Silder, T. K. Uchida, J. L. Hicks, and S. L. Delp, “Simulating ideal assistive devices to reduce the metabolic cost of walking with heavy loads,” PLOS ONE, vol. 12, no. 7, p. e0180320, Jul. 2017, publisher: Public Library of Science. [Online]. Available: https://journals.plos.org/plosone/article?id=10.1371/journal.pone.0180320

[41] N. A. Bianco, P. W. Franks, J. L. Hicks, and S. L. Delp, “Coupled exoskeleton assistance simplifies control and maintains metabolic benefits: a simulation study,” bioRxiv, p. 2021.04.16.440073, Apr. 2021, publisher: Cold Spring Harbor Laboratory Section: New Results. [Online]. Available: https://www.biorxiv.org/content/10.1101/2021.04.16.440073v1

[42] G. M. Bryan, P. W. Franks, S. C. Klein, R. J. Peuchen, and S. H. Collins, “A hip–knee–ankle exoskeleton emulator for studying gait assistance,” The International Journal of Robotics Research, p. 0278364920961452, Nov. 2020, publisher: SAGE Publications Ltd STM. [Online]. Available: https://doi.org/10.1177/0278364920961452

[43] J. Zhang, C. C. Cheah, and S. H. Collins, “Experimental comparison of torque control methods on an ankle exoskeleton during human walking,” in 2015 IEEE International Conference on Robotics and Automation (ICRA). Seattle, WA, USA: IEEE, May 2015, pp. 5584–5589. [Online]. Available: http://ieeexplore.ieee.org/document/7139980/

[44] K. A. Witte, P. Fiers, A. L. Sheets-Singer, and S. H. Collins, “Improving the energy economy of human running with powered and unpowered ankle exoskeleton assistance,” Science Robotics, vol. 5, no. 40, Mar. 2020, publisher: Science Robotics Section: Research Article. [Online]. Available: http://robotics.sciencemag.org/content/5/40/eaay9108

[45] J. R. Koller, D. H. Gates, D. P. Ferris, and C. D. Remy, “‘Body-in-the-Loop’ Optimization of Assistive Robotic Devices: A Validation Study,” in Robotics: Science and Systems, 2016.

[46] N. Hansen, “The CMA Evolution Strategy: A Comparing Review,” in Towards a New Evolutionary Computation: Advances in the Estimation of Distribution Algorithms, ser. Studies in Fuzziness and Soft Computing, J. A. Lozano, P. Larrañaga, I. Inza, and E. Bengoetxea, Eds. Berlin, Heidelberg: Springer, 2006, pp. 75–102.

[47] J. C. Selinger and J. M. Donelan, “Estimating instantaneous energetic cost during non-steady-state gait,” Journal of Applied Physiology, vol. 117, no. 11, pp. 1406–1415, Dec. 2014. [Online]. Available: https://www.physiology.org/doi/10.1152/japplphysiol.00445.2014

[48] J. De Luca, L. Donald Gilmore, M. Kuznetsov, and S. H. Roy, “Filtering the surface EMG signal: Movement artifact and baseline noise contamination,” Journal of Biomechanics, vol. 43, no. 8, pp. 1573–1579, May 2010. [Online]. Available: https://www.sciencedirect.com/science/article/pii/S0021929010000631

[49] A. Winter, The Biomechanics and Motor Control of Human Gait: Normal, Elderly and Pathological, 2nd ed. Waterloo Biomechanics, 1991.

[50] G. J. Bastien, P. A. Willems, B. Schepens, and N. C. Heglund, “Effect of load and speed on the energetic cost of human walking,” European Journal of Applied Physiology, vol. 94, no. 1-2, pp. 76–83, May 2005.

[51] S. D. Prentice, E. N. Hasler, J. J. Groves, and J. S. Frank, “Locomotor adaptations for changes in the slope of the walking surface,” Gait & Posture, vol. 20, no. 3, pp. 255–265, Dec. 2004. [Online]. Available: https://www.sciencedirect.com/science/article/pii/S096663620300167X

[52] R. L. Medrano, G. C. Thomas, and E. Rouse, “Methods for Measuring the Just Noticeable Difference for Variable Stimuli: Implications for Perception of Metabolic Rate with Exoskeleton Assistance,” in 2020 8th IEEE RAS/EMBS International Conference for Biomedical Robotics and Biomechatronics (BioRob), Nov. 2020, pp. 483–490, iSSN: 2155-1782.

[53] R. W. Jackson, C. L. Dembia, S. L. Delp, and S. H. Collins, “Muscle–tendon mechanics explain unexpected effects of exoskeleton assistance on metabolic rate during walking,” Journal of Experimental Biology, vol. 220, no. 11, pp. 2082–2095, Jun. 2017, publisher: The Company of Biologists Ltd Section: Research Article. [Online]. Available: https://jeb.biologists.org/content/220/11/2082

[54] Lee, I. Kang, D. Molinaro, A. Yu, and A. Young, “Real-Time User-Independent Slope Prediction using Deep Learning for Modulation of Robotic Knee Exoskeleton Assistance,” IEEE Robotics and Automation Letters, pp. 1–1, 2021, conference Name: IEEE Robotics and Automation Letters.

[55] P. W. Franks, N. A. Bianco, G. M. Bryan, J. L. Hicks, S. L. Delp, and S. H. Collins, “Testing Simulated Assistance Strategies on a HipKnee-Ankle Exoskeleton: a Case Study,” in 2020 8th IEEE RAS/EMBS International Conference for Biomedical Robotics and Biomechatronics (BioRob), Nov. 2020, pp. 700–707, iSSN: 2155-1782.

[56] S. Song, H. Choi, and S. H. Collins, “Using force data to self-pace an instrumented treadmill and measure self-selected walking speed,” Journal of NeuroEngineering and Rehabilitation, vol. 17, no. 1, p. 68, Jun. 2020. [Online]. Available: https://doi.org/10.1186/s12984-020-00683-5

